# HPV co-infection burden, vaginal microbiota restructuring, and host context are associated with Cervical precancer risk in a racially restricted observational cohort

**DOI:** 10.64898/2026.05.13.724979

**Authors:** Katherine Y. Tossas, Bin Zhu, Katarzyna M. Tyc, Ciara Rhodes, Jerome F. Strauss, Myrna G. Serrano, Gregory A. Buck

**Author notes:** **Correspondence**: Katherine Y. Tossas, PhD, MS; Virginia Commonwealth University. **Co-author emails**.

## Abstract

**Background:** High-risk human papillomavirus (HPV) infection is necessary for cervical carcinogenesis, but HPV detection alone does not distinguish transient infection from lesions at greatest risk of progression. We evaluated whether HPV burden, vaginal microbiota structure, and host-context variables jointly characterize cervical intraepithelial neoplasia grade 3 (CIN3) in a Black/African American and White analytic cohort from the Vaginal Microbiome Health Project (VaMHP), integrating L1-based HPV typing, 16S rRNA vaginal microbiota profiling, and linked clinical metadata.

**Results:** Among 1181 participants, 75 had CIN3. CIN3 was associated with HPV positivity (55/75, 73.3% vs 431/1106, 39.0%; odds ratio [OR] 4.31, 95% CI 2.55-7.29; Fisher exact p = 7.9 x 10^-9) and with multiple HPV infection among HPV-positive participants (35/55, 63.6% vs 176/431, 40.8%; OR 2.54, 95% CI 1.42-4.54; p = 0.0022). HPV communities in CIN3-positive samples showed higher Shannon diversity, greater observed strain richness, higher evenness, and significant beta-diversity separation. In vaginal microbiota analyses, alpha diversity did not differ by CIN3 status, but community composition did, and Lactobacillus crispatus was the only taxon depleted in CIN3 after multiple-testing correction. Race, age, and metronidazole exposure were central nodes in the host-factor network. In predictive modeling, a full integrated model combining metadata, HPV, and vaginal microbiota features (auROC = 0.745) outperformed both HPV + vaginal microbiota (auROC = 0.670) and HPV-only (auROC = 0.440) models.

**Conclusions:** CIN3 in this cohort was associated with coordinated shifts in virologic burden, vaginal community structure, and host social-clinical context. The results support a structure-function interpretation in which loss of Lactobacillus crispatus-dominant states and enrichment of dysbiosis-associated communities define a host-microbiome context that is more permissive to HPV persistence and precancer. These findings move beyond descriptive omics by showing that microbiome and host-context features add nonredundant discriminatory signal beyond HPV-only models.

## Background

Human papillomavirus (HPV) causes virtually all cervical cancers, and persistent infection with oncogenic HPV types is the necessary initiating event for cervical carcinogenesis [1]. Yet HPV detection alone does not reliably identify which individuals will progress from transient infection to high-grade precancer. Most HPV infections resolve spontaneously, and even among those with cervical dysplasia, only a subset progress to cervical intraepithelial neoplasia grade 3 (CIN3), the lesion most closely linked to invasive cervical cancer [1–3]. This discrepancy has driven interest in co-factors that may modulate persistence, immune escape, and epithelial transformation.

One candidate modifier is the vaginal microbiota (VMB). Across multiple studies, Lactobacillus-dominant vaginal communities, especially those enriched for Lactobacillus crispatus, have been associated with favorable genital tract homeostasis, characterized by lower pH, reduced inflammation, and intact mucosal barrier function, whereas diverse anaerobe-rich communities have been linked to HPV persistence, cervical dysplasia, and cancer risk [4–8]. Mechanistically, dysbiotic states may promote epithelial disruption, alter cervicovaginal metabolites, and reshape local immune responses in ways that permit viral persistence and progression [4–7]. These observations suggest that the microbiologic context in which HPV infection occurs may be a critical determinant of cervical cancer risk. Because the VMB is shaped not only by host biology but also by sexual, clinical, psychosocial, and environmental exposures, microbial patterns associated with HPV persistence may also reflect the downstream biologic consequences of structural inequity.

The distribution of these microbial risk states is not random. Cervical precancer and cancer are unequally distributed across racialized and socioeconomically marginalized populations. In the United States, Black women continue to experience disproportionate burden across the cervical cancer continuum, including delayed screening, barriers to follow-up, and worse outcomes after diagnosis [17,18]. Importantly, race should not be interpreted as a biologic essence in this setting; rather, it is a social variable that frequently captures exposure to structurally patterned conditions, including differences in health care access, chronic stress, treatment pathways, and environmental context [17,18]. These same structural conditions may also shape microbial ecology, sexual health exposures, and the likelihood that HPV infection progresses rather than clears.

The Vaginal Microbiome Health Project (VaMHP) provides a rare opportunity to study these processes jointly in a large, racially diverse cohort with paired HPV sequencing, 16S rRNA profiling, and clinical metadata [9]. Using these data, we previously showed that the relationship between the VMB and pre-cervical lesions may differ by race [10]. Building on that foundation, the present study examined whether CIN3 risk is associated with HPV strain burden, HPV community structure, vaginal microbial composition, and host social-clinical context in an integrated analytic framework. We hypothesized that CIN3 would be associated with: (1) greater HPV co-infection burden and distinct HPV community structure; (2) depletion of health-associated Lactobacillus taxa, especially L. crispatus; and (3) a broader network of social and clinical correlates that help contextualize biologic vulnerability.

## Methods

### Study population and design

This analysis used data from the Vaginal Microbiome Health Project (VaMHP), a prospective observational cohort established at Virginia Commonwealth University and affiliated clinics to investigate the vaginal microbiome across gynecologic and reproductive health states [9]. The parent cohort enrolled 2,171 non-pregnant participants between 2009 and 2013. For the present study, we restricted the analytic sample to participants with non-missing CIN3 status, HPV status, and vaginal microbiome profiles, permission for linkage to clinical metadata, and self-identification as Black/African American or White. This restriction was applied because other racial and ethnic groups were sparsely represented in the cohort, precluding stable subgroup analyses. The resulting final analytic cohort included 1,181 participants, of whom 75 were classified as CIN3-positive and 1,106 as CIN3-negative.

Within this cohort, 822 participants (70%) self-identified as Black/African American and 359 (30%) as White. HPV infection was detected in 486 participants (41%). Vaginal bacterial communities were heterogeneous across 10 study-specific community-type categories.

Descriptive characteristics of this final analytic cohort are presented in Supplementary Table 1, and variable-level missingness is summarized in Supplementary Table 2. Clinical and demographic metadata included CIN3 status, self-reported race, age, selected medical history, medication exposures, behavioral variables, and socioeconomic indicators. The primary outcome for the present analysis was CIN3.

### HPV amplification, sequencing, and typing

HPV status and type were determined using a two-step PCR protocol targeting a 452-bp region of the L1 gene, corresponding to nucleotide positions 946–1397 of the HPV16 L1 gene reference sequence. In the first step, the target region was amplified using consensus L1-specific primers carrying partial Illumina adapter sequences: forward primer HPV-01F (5’-TCGTCGGCAGCGTCAGATGTGTATAAGAGACAGGCACAGGGMCAHAAYAATGG-3’) and reverse primer HPV-6R (5’-GTCTCGTGGGCTCGGAGATGTGTATAAGAGACAGNCGTCCCAAAGGRAWYTGATC- 3’). Reactions were performed in a 26 µL volume containing approximately 20 ng of DNA template, 0.2 µM of each primer, 10 mM dNTPs, 3.75 U of AmpliTaq DNA Polymerase (Thermo Fisher), 1.5 µL of 25 mM MgCl, and 2.5 µL of the supplied 10X buffer.

Thermocycling conditions followed Gravitt et al. (2000): initial denaturation at 95°C for 9 minutes, 40 cycles of 95°C for 60 seconds, 55°C for 60 seconds, and 72°C for 60 seconds, with a final extension at 72°C for 5 minutes. First-step amplicons were purified using 0.8X Agencourt AMPure XP beads (Beckman Coulter Genomics) per the manufacturer’s protocol and resuspended in 25 µL of Teknova DNA Suspension Buffer. In the second step, sample-specific barcodes and full Illumina sequencing adapters were attached. Reactions contained 5 µL of first-step product diluted 1:10 as template, 0.05 µM Nextera indexing primers, and 12.5 µL of Q5 High-Fidelity 2X Master Mix (New England Biolabs) in a total volume of 25 µL. Thermocycling proceeded with an initial denaturation at 98°C for 5 minutes, 12 cycles of 95°C for 30 seconds, 62°C for 60 seconds, and 72°C for 60 seconds, and a final extension at 72°C for 10 minutes.

Second-step amplicons were purified using 0.8X AMPure XP beads. DNA concentration was quantified using the QuantiFluor dsDNA System (Promega) on a BioTek Synergy HT plate reader, and fragment size distribution was confirmed on an Agilent TapeStation. Indexed amplicons were pooled at equimolar concentrations and sequenced on an Illumina MiSeq platform using a 600-bp paired-end configuration.

Sequencing reads were demultiplexed using sample-specific barcodes, adapter sequences were removed, and human reads were filtered prior to HPV alignment. To exclude human reads, sequences were first aligned against the human reference genome GRCh38.v41 using NextGenMap (version 0.5.5) and reads aligning with more than 95% identity over at least 75% of their length were removed. The remaining non-human reads were merged and quality-filtered using MeFiT before downstream analysis [15], then mapped against a reference collection of HPV L1 sequences representing 229 officially established HPV types sourced from the Karolinska Institute. Reads mapping to HPV L1 sequences were required to meet a minimum of 90% identity over at least 75% of their length. An HPV type was considered present when at least 25 reads mapped to that type and those reads represented more than 5% of non-human reads in the sample.

### Vaginal microbiota profiling

Vaginal bacterial communities were characterized by 16S rRNA gene sequencing of the V1-V3 regions using the VaMHP barcoded PCR workflow described previously [9]. Raw sequence reads were demultiplexed using sample-specific barcodes, adapter sequences were trimmed, human reads were filtered out, and overlapping paired-end reads were merged and quality-filtered using MeFiT with a maximum expected error (meep) cutoff of 1.0. High-quality reads were then taxonomically assigned using the STIRRUPS pipeline and reference database [16].

Because reference sequences in the STIRRUPS database are assigned to species-level clusters based on ≥97% identity across the V1–V3 region [16], closely related taxa whose sequences fall within this threshold are represented as a single cluster rather than as discrete species. For example, the *Peptoniphilus*_cluster encompasses sequences assigned to *P. vaginalis* (NR_179440.1), *P. indolicus* (NR_115509.1 and LN871839), and *P. harei* (LRQE01000014), which are insufficiently divergent in the V1–V3 region to permit reliable species-level discrimination from short amplicon reads alone. It should be noted, however, that additional sequences for *P. indolicus* and *P. harei* are available in NCBI that do not classify within this cluster, indicating that cluster membership is sequence-specific rather than species-wide.

Consequently, species-level designations reported throughout this manuscript should be interpreted as cluster-level assignments where applicable, with the caveat that cluster composition may not fully capture the true diversity of the named species. Samples were retained only if they contained more than 5,000 total reads. Prior to differential analyses, 16S rRNA feature tables were pre-filtered to retain taxa with relative abundances exceeding 0.1% in more than 5% of samples or at least 0.01% in more than 15% of samples, in order to reduce sparsity and limit unstable inference driven by extremely rare taxa. Negative and positive controls were incorporated into the laboratory workflow, including extraction blanks and PCR no-template controls, to monitor potential contamination and amplification artifacts.

### Statistical analysis

Associations between metadata variables were assessed according to variable type using Spearman correlation, point-biserial correlation, or chi-square testing, with Benjamini-Hochberg false-discovery-rate correction applied where specified. HPV strain-level associations with CIN3 were tested using Fisher exact tests, followed by false-discovery-rate correction. The overall type-distribution comparison shown in Figure 1C was assessed using Fisher’s exact test with Monte Carlo simulation. HPV alpha diversity was summarized using the Shannon index, observed strain richness, and evenness. Vaginal microbiota alpha diversity was summarized using the Shannon index. Beta diversity for both HPV and vaginal microbiota communities was evaluated using Bray-Curtis dissimilarity, nonmetric multidimensional scaling, and PERMANOVA implemented in vegan [12]. Differential abundance analyses for vaginal taxa were performed with ALDEx2 [13].

**Figure 1.**
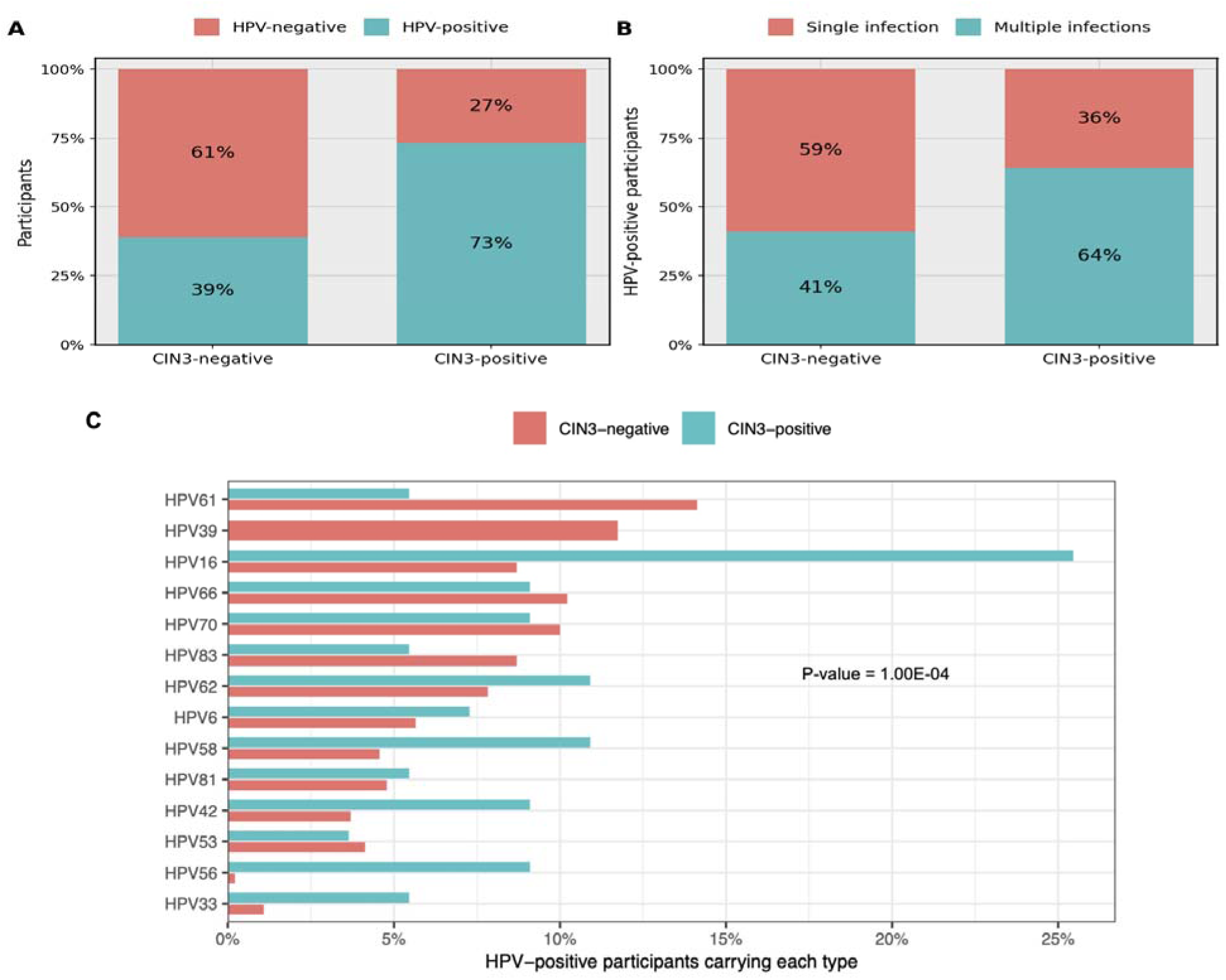
HPV prevalence and type distribution by CIN3 status. (A) Percentage of HPV-negative and HPV-positive participants in CIN3-negative and CIN3-positive groups. (B) Among HPV-positive participants, percentage with single-type versus multiple-type infections by CIN3 status. (C) Distribution of selected HPV types among HPV-positive CIN3-negative and CIN3-positive participants.

To investigate multilevel interrelationships among host variables, significantly associated metadata dyads were used to build an unweighted network in Gephi, with node size determined by betweenness centrality and clusters identified through modularity analysis [14]. The host-factor network included 29 demographic, behavioral, clinical, and microbial-context variables drawn from the VaMHP questionnaire and linked study metadata. These included race, age, education, employment, income, marital status, smoking, alcohol use frequency, lifetime and past-year number of sexual partners, douching, recurrent yeast infection, history of bacterial vaginosis, recent medication exposures (metronidazole, penicillin, tetracyclines, macrolides, quinolones, nitrofurantoin, and insulin), vegetarian and vegan diet indicators, vaginal microbiome type, cancer history/site, and CIN3 status. For predictive modeling, three random forest models were evaluated to compare the incremental contribution of different data domains: an HPV-only model, an HPV + vaginal microbiota (VMB) model, and a full integrated model including metadata, HPV, and VMB predictors. Ordinal variables were transformed and categorical variables were one-hot encoded before model fitting. Metadata variables with missing values were imputed using random forest proximity imputation with the rfImpute function in R before model fitting; the final selected predictors entering the full integrated model had no remaining missing values. No explicit class-imbalance correction was applied. Model performance was summarized by area under the receiver operating characteristic curve (auROC), and variable importance in the full integrated model was ranked by mean decrease in Gini.

## Results

### Overview of the analytic cohort

The analytic cohort comprised 1,181 non-pregnant Black/African American and White participants with linked clinical, HPV, and vaginal microbiome data. CIN3 was identified in 75 participants (6.4%), while 1,106 participants were classified as CIN3-negative. Within this cohort, 822 participants (70%) self-identified as Black/African American and 359 (30%) as White, and HPV infection was detected in 486 participants (41%). Vaginal bacterial communities were heterogeneous across 10 study-specific community-type categories. Descriptive characteristics of the analytic cohort are provided in Supplementary Table 1, and variable-level missingness is summarized in Supplementary Table 2.

### HPV burden and strain-level associations with CIN3

HPV infection was substantially more common among participants with CIN3 than among those without CIN3 (55/75, 73% vs 431/1,106, 39%; Figure 1A). HPV positivity itself was strongly associated with CIN3 (odds ratio [OR] = 4.31, 95% confidence interval [CI]: 2.55–7.29, Fisher exact p = 7.9 × 10^-9). Among HPV-positive participants, single infections predominated overall (275/486, 56.6%), but multiple infections were enriched in the CIN3-positive group (35/55, 63.6% vs 176/431, 40.8%; Figure 1B), corresponding to an OR of 2.54 with 95% CI: 1.42–4.54 for multiple infection among HPV-positive participants with CIN3 (Fisher exact p = 0.0022).

Type distributions also appeared to differ by disease status (Figure 1C), indicating that CIN3 tracked both with HPV presence and with the composition of mixed-type infections.

HPV community structure also differed by disease status (Figure 2). Relative-composition plots suggested broader multi-type HPV profiles among CIN3-positive participants (Figure 2A). Across alpha-diversity metrics, CIN3-positive samples showed higher Shannon diversity (p = 0.0025), greater observed strain richness (p = 0.0147), and higher evenness (p = 0.0183) (Figure 2B-D). Beta diversity also differed significantly by disease status (PERMANOVA p = 0.001; Figure 2E). At the strain level, four HPV types remained significantly associated with CIN3 positivity after Benjamini-Hochberg correction: HPV16 (FDR = 1.21 x 10^-9), HPV56 (FDR = 5.03 x 10^-11), HPV33 (FDR = 0.024), and HPV42 (FDR = 0.024) (Table 1). HPV16 and HPV56 provided the clearest canonical oncogenic signal, whereas the HPV42 association should be interpreted cautiously because it may reflect the ecology of mixed infections rather than independent carcinogenicity.

**Figure 2.**
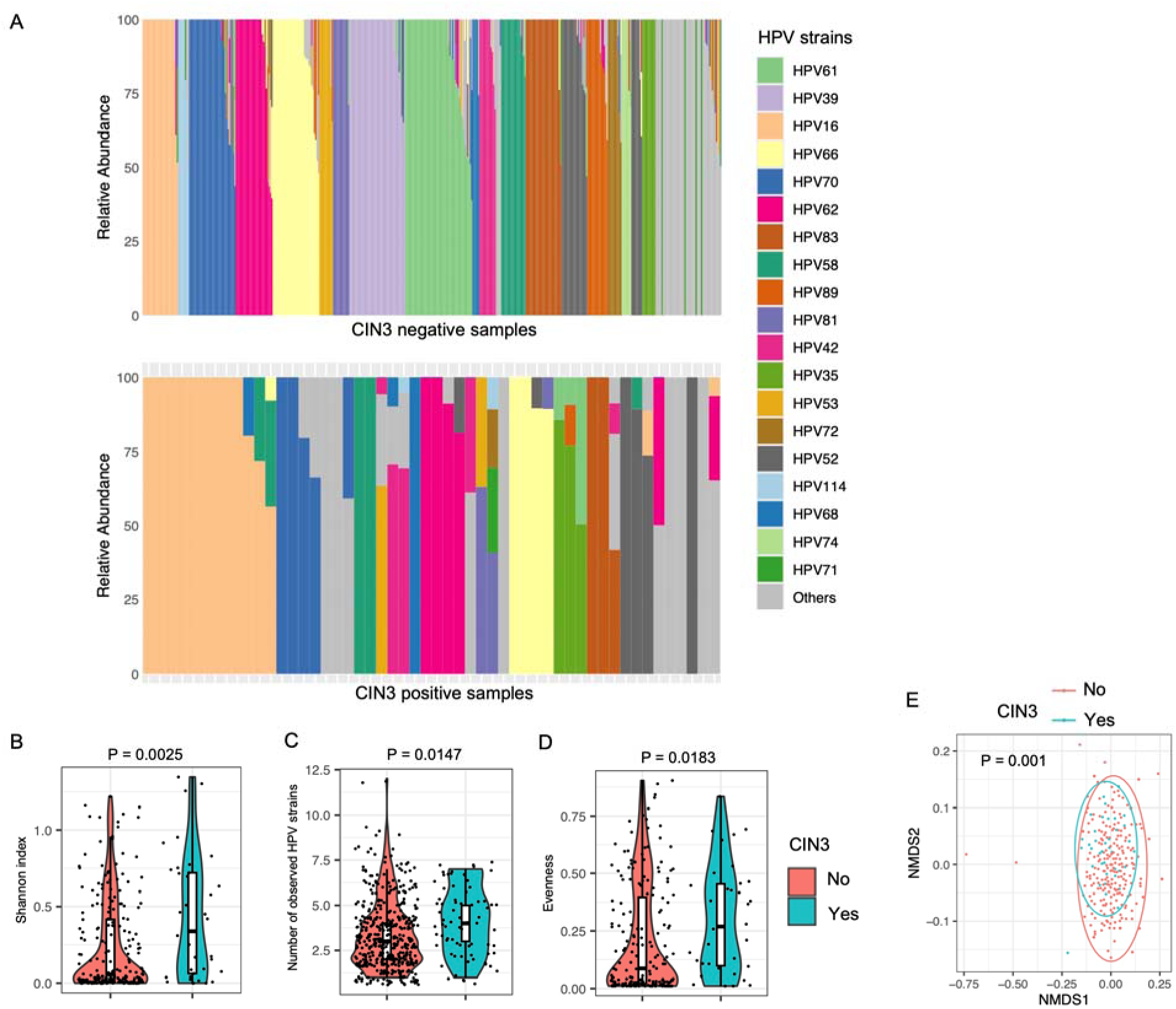
HPV community structure in relation to CIN3. (A) Stacked bar plots of HPV strain composition among HPV-positive samples, shown separately for CIN3-negative and CIN3-positive participants. (B-D) Shannon diversity, observed HPV strain richness, and evenness by CIN3 status, with p-values from Mann-Whitney tests. (E) NMDS plot of HPV beta diversity by CIN3 status, with significance assessed by Adonis/PERMANOVA.

**Table 1.** HPV types significantly associated with CIN3 after false-discovery-rate correction.

| HPV type | Direction of association | FDR-adjusted p value |
| --- | --- | --- |
| HPV16 | Enriched in CIN3-positive participants | 1.21 x 10<sup>-9</sup> |
| HPV56 | Enriched in CIN3-positive participants | 5.03 x 10<sup>-11</sup> |
| HPV33 | Enriched in CIN3-positive participants | 0.024 |
| HPV42 | Enriched in CIN3-positive participants | 0.024 |

### Race-stratified HPV patterns

In descriptive race-stratified summaries (Figure 3), no detectable HPV remained the most common category among CIN3-negative participants in both racial groups, accounting for 67% of White participants and 58% of Black/African American participants. Among CIN3-positive participants, the distribution shifted toward greater HPV burden in both groups. Within White participants, the proportion with no detectable HPV decreased to 22%, while the proportions with single and multiple infections increased to 44% and 33%, respectively. Within Black/African American participants, the proportion with no detectable HPV decreased to 27%, while multiple infection rose from 17% in CIN3-negative participants to 48% in CIN3-positive participants and exceeded the proportion with single infection (24%). These patterns suggest that the link between CIN3 and greater co-infection burden was present in both groups but may have been more strongly driven by multiple infections among Black/African American participants.

**Figure 3.**
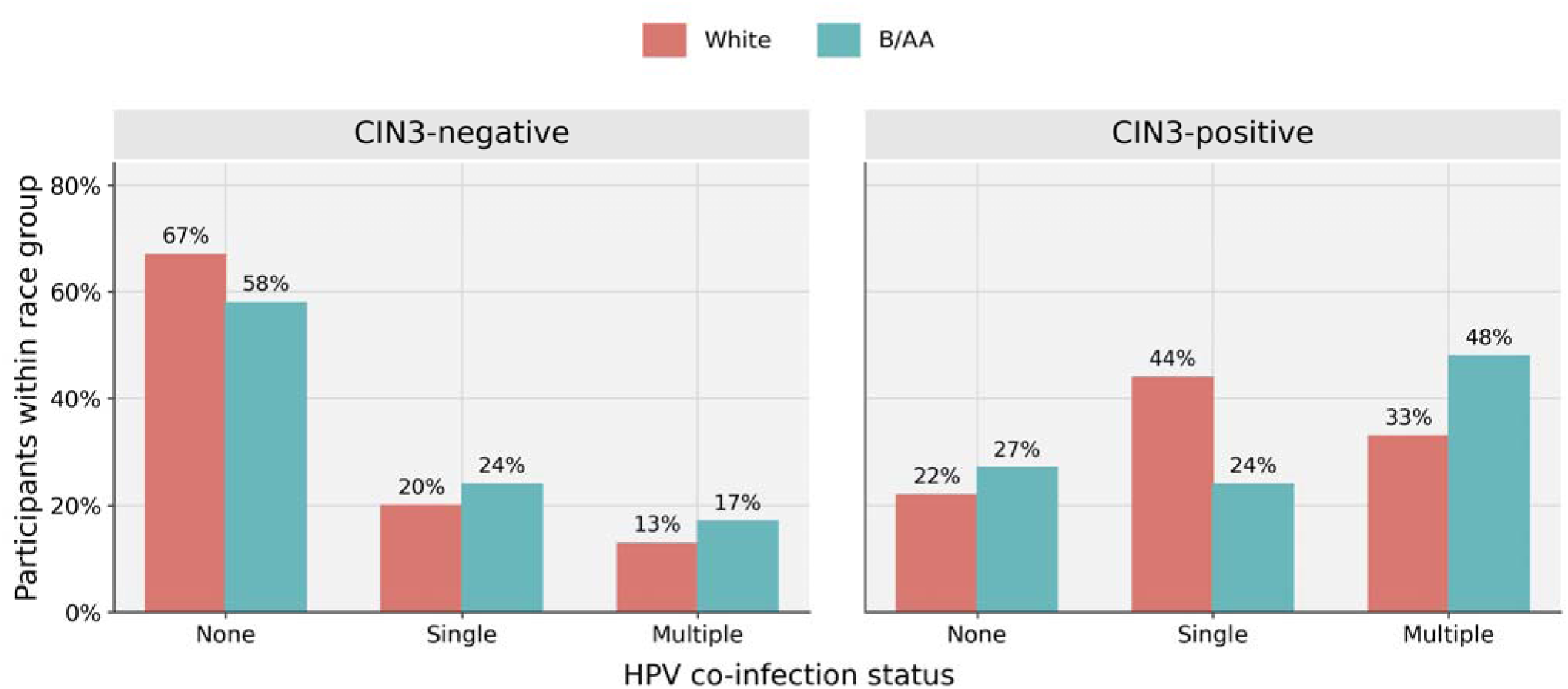
Descriptive race-stratified distribution of HPV infection burden by CIN3 status. Bars show the percentage of White and Black/African American participants within each CIN3 stratum who had no detectable HPV, a single-type HPV infection, or multiple-type HPV infection. Percentages are calculated within race group and CIN3 stratum. This figure is intended as a descriptive race-stratified summary and is not presented as formal subgroup inference.

Because subgroup counts, particularly among White CIN3-positive participants, were limited, these comparisons are presented descriptively and should not be interpreted as formal race-stratified inference.

Taken together, these analyses indicate that CIN3 in this cohort tracked more closely with virologic burden - especially multi-type infection and higher HPV community diversity - than with HPV positivity alone. Any apparent race-specific gradients should nevertheless be interpreted cautiously because subgroup sizes, particularly among White participants with CIN3, were small.

### Vaginal microbiota correlates of CIN3

The vaginal microbiota showed compositional differences by CIN3 status (Figure 4). Overall alpha diversity did not differ significantly (Shannon p = 0.41; Figure 4B). In contrast, beta diversity analyses demonstrated a significant global association between VMB composition and CIN3 (PERMANOVA p = 0.004; Figure 4C), indicating that the overall structure of vaginal bacterial communities differed between CIN3-positive and CIN3-negative participants. Community-composition plots were consistent with greater depletion of Lactobacillus-dominant states and broader interindividual heterogeneity among CIN3-positive samples (Figure 4A).

**Figure 4.**
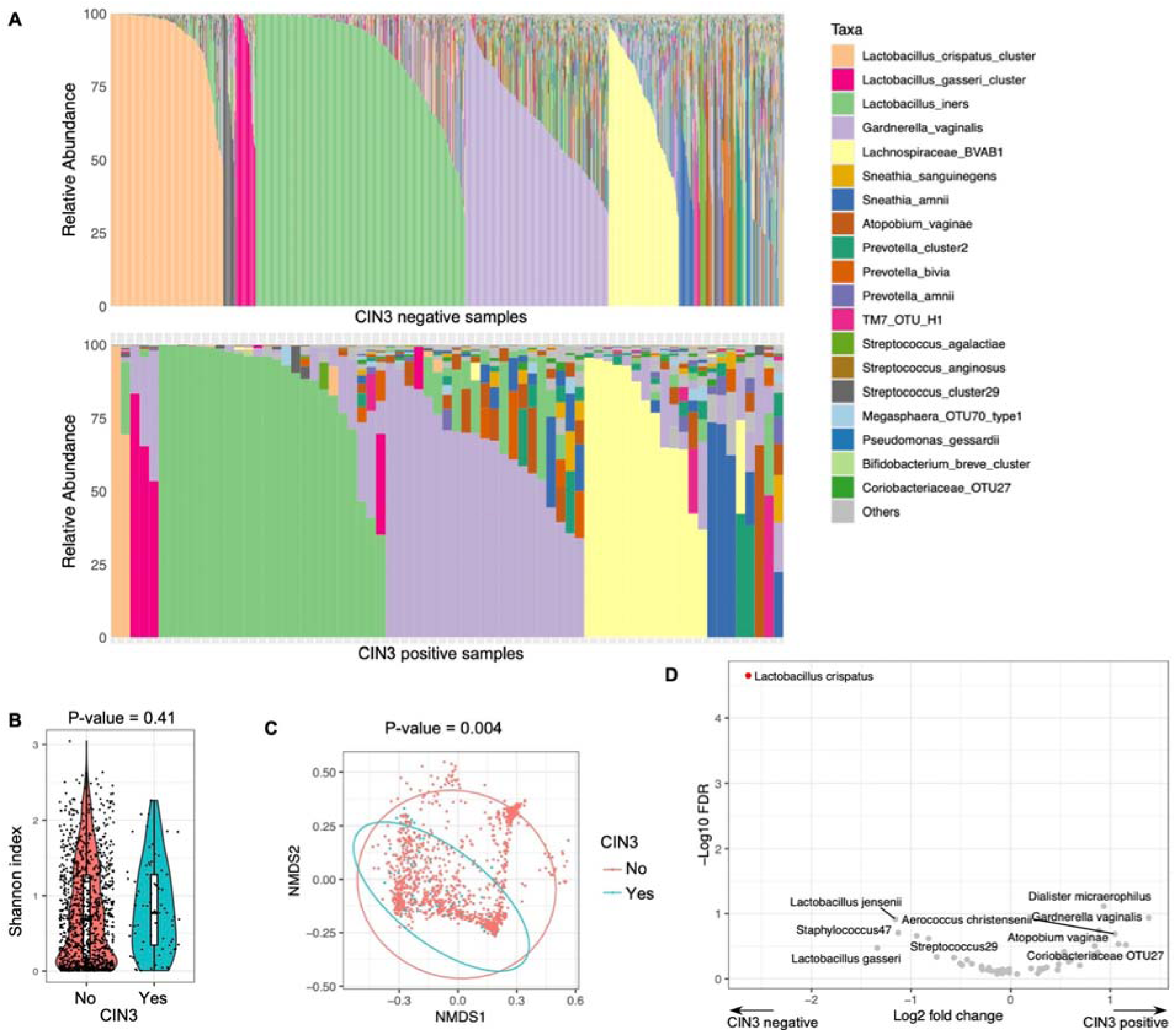
Vaginal microbiota profiles by CIN3 status. (A) Stacked bar plots of vaginal community composition in CIN3-negative and CIN3-positive samples. (B) Shannon diversity by CIN3 status. (C) NMDS plot of vaginal microbiota beta diversity by CIN3 status, with significance assessed by Adonis/PERMANOVA. (D) Differential-abundance plot of vaginal taxa associated with CIN3; positive log2 fold changes indicate relative enrichment in CIN3-positive samples and negative values indicate relative depletion. Lactobacillus crispatus was the only taxon meeting the false-discovery-rate threshold.

At the taxonomic level, *L. crispatus* was the only vaginal taxon that remained significantly depleted in CIN3-positive samples after multiple-testing correction (Figure 4D). Other *Lactobacillus* species, including *L. jensenii* (P = 0.296, FDR = 0.716) and *L. gasser*i (P = 0.751, FDR = 0.909), were also lower in CIN3-positive participants but did not meet the multiple-testing threshold for statistical significance. In contrast, anaerobe-associated taxa, including *Gardnerella vaginalis, Fannyhessea vaginae* (formerly classified as *Atopobium vaginae*), and *Dialister micraerophilus*, were enriched in CIN3-positive samples. Race-stratified analyses suggested that the magnitude of these VMB-CIN3 contrasts may differ across groups, although those subgroup comparisons were underpowered.

### Host-factor network and multivariable prediction

To situate microbial and virologic findings within a broader host context, the study evaluated pairwise associations across 29 demographic and clinical variables and constructed a host-factor network from significant associations (Figure 5). In that network, race (bw) emerged as a central hub linking multiple sociodemographic and clinical variables and had the highest betweenness centrality score in the network (43.04). Metronidazole exposure (36.38) and age (30.38) were also among the most central nodes, indicating that CIN3 was embedded within a broader structure of correlated social and clinical factors. The full set of variables included in the network is provided in the Methods and node-level centrality statistics are available in Supplementary Dataset 1. CIN3 was directly connected to race, marital status, employment, education, yearly alcohol consumption, income, and a cancer-diagnosis variable, placing disease status within a broader module of structurally and behaviorally patterned factors.

**Figure 5.**
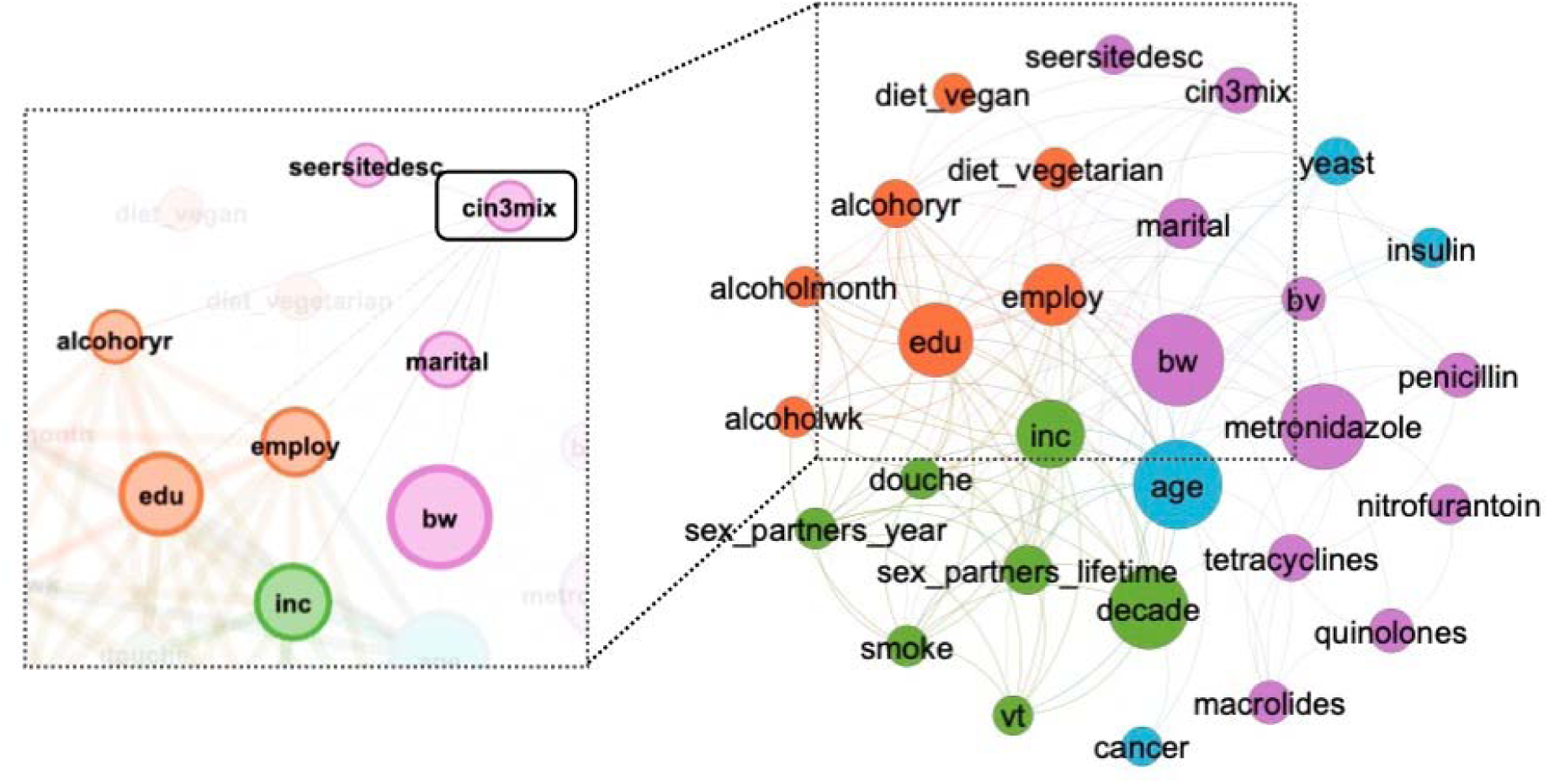
Host-factor association network. Nodes represent host variables, node size corresponds to betweenness centrality, and colors indicate modularity class. Edges represent statistically significant pairwise associations among variables. The left panel highlights the neighborhood of CIN3-associated variables, whereas the right panel shows the broader network structure.

A multivariable random forest model integrating metadata, HPV, and vaginal microbiota predictors achieved an auROC of 0.745 (Figure 6A), indicating moderate discrimination. This full integrated model outperformed both the HPV + VMB model (auROC = 0.670) and the HPV-only model (auROC = 0.440), supporting the added predictive value of combining social-clinical context with virologic and microbial features. Variable-importance rankings from the full integrated model indicated that vaginal microbial features were among the most influential predictors, with *G. vaginalis, L. crispatus*, and the *Peptoniphilus* cluster ranking above most individual HPV types (Figure 6B). HPV co-infection burden (HPV_number) was also a top predictor, followed by income, HPV56, HPV16, education, employment, and marital status. The correlation heatmap suggested that microbial taxa, HPV burden, and social-clinical variables formed partially overlapping but nonredundant clusters (Figure 6C), supporting a multilevel model in which CIN3 is associated with the convergence of viral burden, vaginal dysbiosis, and social-clinical context rather than any single factor in isolation.

**Figure 6.**
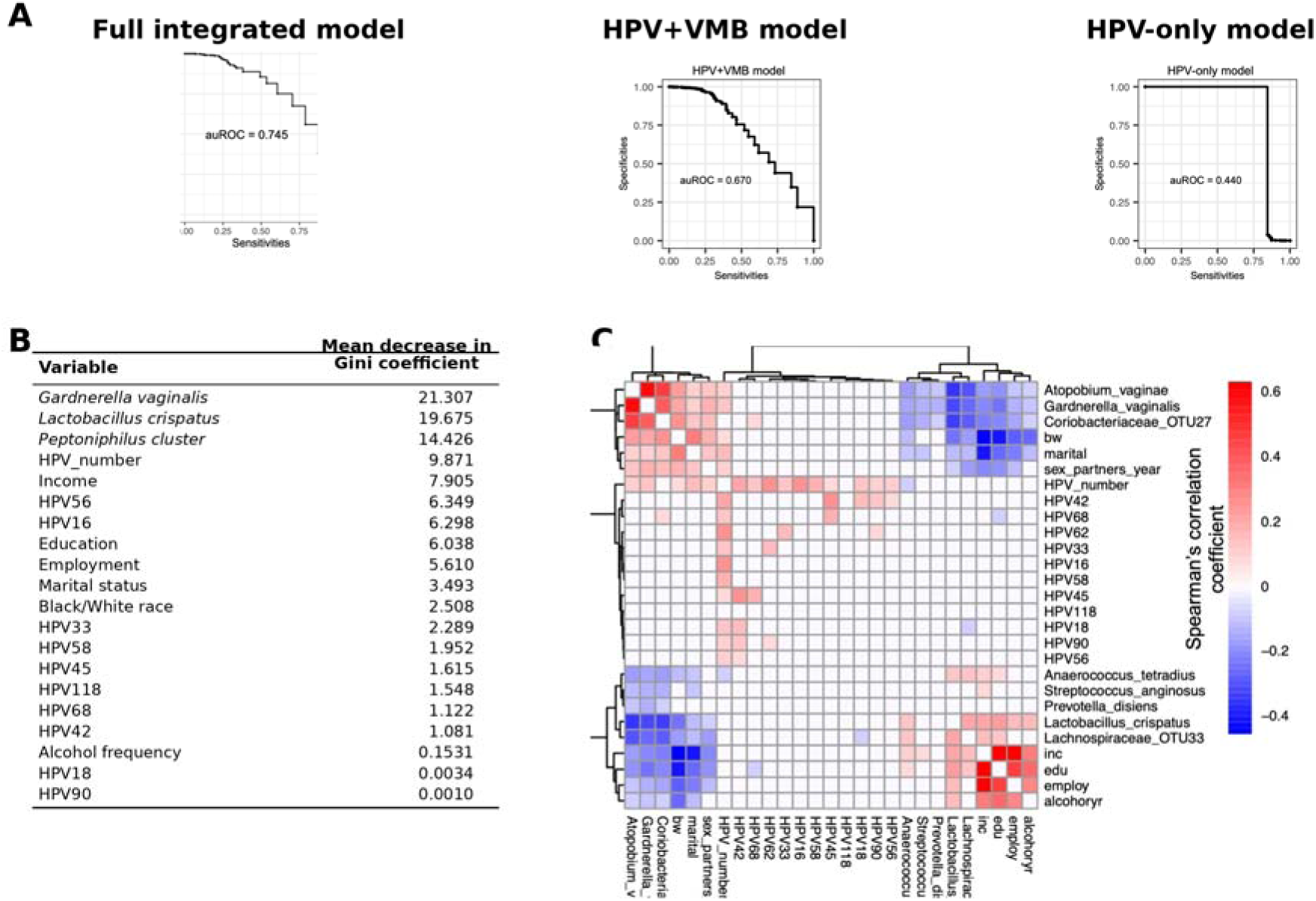
Integrative prediction of CIN3. (A) Comparison of random forest model performance across an HPV-only model (auROC = 0.440), an HPV + vaginal microbiota (VMB) model (auROC = 0.670), and the full integrated metadata + HPV + VMB model (auROC = 0.745). (B) Variable-importance ranking for the full integrated model based on mean decrease in Gini. (C) Heatmap of Spearman correlations among selected predictors used in the full integrated model. The comparator models were nested such that the HPV-only model included virologic predictors alone, the HPV + VMB model included virologic and microbial predictors, and the full model additionally incorporated metadata variables.

## DISCUSSION

In this study, we asked whether cervical precancer is better characterized by integrated virologic, microbial, and host-context profiles than by HPV detection alone. Four findings stand out. First, CIN3 was strongly associated with virologic burden, including both HPV positivity and co-infection with multiple HPV types. Second, CIN3-positive samples harbored more diverse HPV communities, indicating that disease status tracked with community structure as well as with specific HPV types. Third, CIN3 was associated with altered vaginal microbial composition, most notably depletion of *Lactobacillus crispatus*. Fourth, host social-clinical context, summarized in the network analysis and multivariable model, remained intertwined with virologic and microbial patterns. Together, these findings support a host-microbiome interpretation of cervical precancer risk in which pathogen burden, vaginal ecosystem structure, and socially patterned context converge.

The virologic findings are biologically plausible and consistent with prior literature. HPV16 and HPV33 are established high-risk types, and persistent infection with oncogenic strains is central to progression from infection to high-grade precancer [1,3]. The identification of HPV56 in the current analysis aligns with work suggesting that a broader range of high-risk or probably high-risk types can contribute to clinically meaningful neoplasia. More informative than any single type, however, was the overall structure of HPV communities: CIN3-positive samples showed greater strain richness, higher evenness, higher Shannon diversity, and greater co-infection burden. These patterns do not prove that multi-type infection is independently causal, but they are consistent with a virologic milieu marked by repeated exposure, impaired clearance, or both. In that sense, the relevant signal may be community-level virologic structure rather than presence or absence of a single genotype.

The microbiome findings advance the study beyond descriptive 16S surveys because they identify a specific ecological configuration associated with CIN3 and link that structure to plausible host interaction pathways. Alpha diversity did not differ by CIN3 status, whereas beta diversity and differential abundance did, indicating that the relevant signal was not generalized diversity but reorganization of community structure. In particular, depletion of *L. crispatus* is clinically meaningful, not merely descriptive. *L. crispatus*-dominant communities are functionally associated with lactic acid production, maintenance of low vaginal pH, support of epithelial barrier integrity, and reduced pro-inflammatory tone [4–8,11]. Loss of this state, together with enrichment of *Gardnerella, Fannyhessea*, and *Dialister*, is therefore consistent with a transition toward a community architecture more permissive to HPV persistence and mucosal dysregulation. Even without direct metabolomic or cytokine measurements in the present study, the observed structure is mechanistically coherent with known host-microbiome interaction pathways.

The third-ranked microbial predictor corresponded to the *Peptoniphilus* cluster, a cluster-level STIRRUPS assignment that may encompass multiple closely related taxa. Although the variable-importance display in Figure 6B labels this feature as *Peptoniphilus* cluster, under the STIRRUPS framework it should be interpreted conservatively as a cluster-level signal rather than definitive evidence for a single species-specific effect. Given the cross-sectional design and cluster-level taxonomic resolution, whether this association reflects an independent contribution to CIN3 risk or co-occurrence with other dysbiosis-associated taxa remains unresolved. More broadly, the fact that microbial features ranked above most individual HPV types in the integrated model argues against treating the microbiome as a passive correlate; instead, the data support the view that community structure carries biologically and clinically relevant information not captured by HPV typing alone.

The host-factor network extends this interpretation by showing that microbial and virologic signals were embedded in a broader host context. Race, age, and metronidazole exposure were the most central nodes in the network, and CIN3 clustered with income, employment, education, marital status, and alcohol use. This network is not causal, but it demonstrates that the microbial states associated with CIN3 do not arise in isolation. Rather, they are interwoven with exposures that shape sexual health, medication use, access to care, and the likelihood that dysbiotic states are acquired, treated, or recur. The findings therefore support a biosocial framing in which race is not interpreted as an intrinsic biologic determinant, but as an index of structurally patterned exposures that can become embedded in clinical, immunologic, and microbial pathways [17,18]. The predictive modeling results reinforce that point. The full metadata + HPV + vaginal microbiota model outperformed both the HPV-only model and the HPV + vaginal microbiota model, showing that host-context variables contributed nonredundant signal beyond virology and microbiota alone. Importantly, the model did not simply recover established carcinogenic HPV types; it prioritized *G. vaginalis, L. crispatus*, the *Peptoniphilus* cluster, HPV co-infection burden, and socioeconomic variables. This benchmarked comparison strengthens the translational relevance of the microbiome findings because it demonstrates that ecological features of the vaginal community are informative for risk discrimination, not merely associated with it. In this respect, the work moves beyond descriptive omics toward a more clinically interpretable host-microbiome framework.

This study has several strengths. It leverages one of the largest racially diverse cohorts with paired HPV and vaginal microbiota sequencing, allowing simultaneous analysis of virologic and microbial features in relation to CIN3 [9,10]. The availability of host metadata enabled a broader systems-level analysis than is typical in microbiome studies. Because VMB sampling preceded later ascertainment of CIN3 through linked records, the study also offers some temporal separation between microbial exposure and clinical outcome, even though temporality for HPV acquisition and persistence remains uncertain.

Several limitations deserve emphasis. First, the analysis is observational and cannot establish causality. Second, although VMB sampling preceded later CIN3 ascertainment, the timing of HPV acquisition, persistence, clearance, and lesion progression cannot be fully resolved from the available data. Third, some subgroup analyses, especially among White CIN3-positive participants, were limited by small numbers. Fourth, race and socioeconomic indicators are imperfect proxies for complex exposures shaped by structural inequity, and residual confounding is likely. Fifth, the study did not include direct inflammatory, immunologic, metabolomic, or transcriptomic measurements that could more directly link community structure to function.

Finally, the predictive modeling analysis was conducted in the setting of class imbalance and has not yet been externally validated; therefore, although the integrated model outperformed reduced-input comparator models, its translational implications should still be interpreted cautiously until replicated in independent cohorts.

Despite these limitations, the study advances the genital microbiome field in two important ways. Conceptually, it links a specific ecological pattern - loss of *L. crispatus*-dominant states, enrichment of dysbiosis-associated anaerobes, and greater interindividual heterogeneity - to a clinically meaningful precancer endpoint rather than to nonspecific gynecologic symptoms alone. Translationally, it shows that microbiome and host-context information improve discrimination beyond HPV-only models, nominating community composition as a plausible adjunct to HPV-based triage. Future work should pair longitudinal HPV measurements with metabolomic, immunologic, and transcriptomic readouts to test whether these community configurations act through acidity, biofilm formation, barrier disruption, inflammation, or combinations thereof, and to determine whether integrated models improve clinically useful risk stratification.

## CONCLUSIONS

CIN3 in the VaMHP cohort was associated with higher HPV burden, altered HPV community structure, compositional shifts in the vaginal microbiota, and a broader social-clinical context in which race functioned as a central correlational node. The strongest microbial signal was depletion of *L. crispatus*, while the strongest virologic signals were multi-type HPV infection and enrichment of several CIN3-associated HPV types, particularly HPV16 and HPV56. In the integrated prediction model, microbial features ranked above most individual HPV types, and the full metadata + HPV + vaginal microbiota model outperformed both HPV-only and HPV + vaginal microbiota models, reinforcing the view that CIN3 risk is multilevel rather than type-deterministic. Together, these findings argue for moving from type-only paradigms toward ecology-informed models of cervical precancer risk.

## Supporting information

STROBE Checklist

## List of abbreviations

CIN3: cervical intraepithelial neoplasia grade 3
HPV: human papillomavirus
VMB: vaginal microbiota
VaMHP: Vaginal Microbiome Health Project
NMDS: non-metric multidimensional scaling
OR: odds ratio
CI: confidence interval
FDR: false-discovery rate
PERMANOVA: permutational multivariate analysis of variance
auROC: area under the receiver operating characteristic curve

## Declarations

### Ethics approval and consent to participate

The parent Vaginal Microbiome Health Project and related linked analyses were reviewed and approved by the Virginia Commonwealth University Institutional Review Board (IRB protocol HM20010007) and collaborating clinical sites, and all participants provided written informed consent before enrollment. All data used in the present analysis were de-identified prior to analysis.

### Consent for publication

Not applicable.

## Availability of data and materials

Raw 16S rRNA gene sequencing data from the Vaginal Human Microbiome Project (VaHMP) are publicly available through the NCBI Sequence Read Archive under BioProject PRJNA46877 and SRA study SRP002462. HPV typing in the present study was performed using a custom reference collection of HPV L1 sequences representing 229 officially established HPV types sourced from the Karolinska Institute. Processed analytic files, Supplementary Tables 1 and 2, node-level centrality statistics, differential-abundance output underlying Figure 4D, and the source-data workbook are provided as additional files with this submission. Analysis scripts used for data processing, diversity analyses, differential abundance, network construction, and predictive modeling are available from the corresponding author on reasonable request. A public code repository will be added prior to final publication if required by the journal.

## Authors’ contributions

KYT conceptualized the study, acquired funding, supervised the work, and drafted the manuscript. BZ performed formal analyses, contributed software, and assisted with drafting and revision. KMT performed formal analyses, contributed software, and assisted with drafting and revision. CR contributed to drafting and revision. JFSIII supervised the work and contributed to revision. MGS contributed to conceptualization, data curation, methodology, and revision. GAB contributed to conceptualization, data curation, methodology, funding acquisition, resources, software, supervision, and revision. All authors read and approved the final manuscript.

## Acknowledgements

We thank the participants in the Vaginal Microbiome Health Project and the clinical and laboratory teams whose work made this analysis possible.

## Additional files

**Additional file 1.** File format: DOCX. Title: Supplementary Table 1. Description: Participant characteristics in the final Black/African American and White analytic cohort overall and by CIN3 status.

**Additional file 2.** File format: DOCX. Title: Supplementary Table 2. Description: Variable-level missingness across candidate predictors in the final analytic cohort.

**Additional file 3.** File format: XLSX. Title: Supplementary Dataset 1. Description: Node-level centrality statistics for the host-factor association network.

**Additional file 4.** File format: XLSX. Title: Supplementary Dataset 2. Description: Differential-abundance results underlying Figure 4D.

**Additional file 5.** File format: DOCX. Title: STROBE checklist. Description: STROBE checklist for this observational cohort study.

**Additional file 6.** File format: XLSX. Title: Source data workbook. Description: Source data underlying key summary figures and Table 1.

## References

1. Schiffman M, Castle PE, Jeronimo J, Rodriguez AC, Wacholder S. Human papillomavirus and cervical cancer. Lancet. 2007;370:890–907.

2. Loopik DL, Bentley HA, Eijgenraam MN, Inthout J, Bekkers RLM, Bentley JR. The natural history of cervical intraepithelial neoplasia grades 1, 2, and 3: a systematic review and meta-analysis. J Low Genit Tract Dis. 2021;25:221–231.

3. Nedjai B, Reuter C, Ahmad A, Banwait R, Warman R, Carton J, et al. Molecular progression to cervical precancer, epigenetic switch or sequential model? Int J Cancer. 2018;143:1720–1730.

4. Sharifian K, Shoja Z, Jalilvand S. The interplay between human papillomavirus and vaginal microbiota in cervical cancer development. Virol J. 2023;20:73.

5. Alimena S, Davis J, Fichorova RN, Feldman S. The vaginal microbiome: a complex milieu affecting risk of human papillomavirus persistence and cervical cancer. Curr Probl Cancer. 2022;46:100877.

6. Dabee S, Passmore JAS, Heffron R, Jaspan HB. The complex link between the female genital microbiota, genital infections, and inflammation. Infect Immun. 2021;89:e00487–20.

7. Anahtar MN, Gootenberg DB, Mitchell CM, Kwon DS. Cervicovaginal microbiota and reproductive health: the virtue of simplicity. Cell Host Microbe. 2018;23:159–168.

8. Mitra A, MacIntyre DA, Ntritsos G, Smith A, Tsilidis KK, Marchesi JR, et al. The vaginal microbiota associates with the regression of untreated cervical intraepithelial neoplasia 2 lesions. Nat Commun. 2020;11:1999.

9. Fettweis JM, Brooks JP, Serrano MG, Sheth NU, Girerd PH, Edwards DJ, et al. Differences in vaginal microbiome in African American women versus women of European ancestry. Microbiology (Reading). 2014;160:2272–2282.

10. Tossas KY, Zhu B, Perera RA, Serrano MG, Sullivan S, Sayeed S, et al. Does the vaginal microbiome operate differently by race to influence risk of precervical cancer? J Womens Health (Larchmt). 2023;32:553–560.

11. Zhu B, Tao Z, Edupuganti L, Serrano MG, Buck GA. Roles of the microbiota of the female reproductive tract in gynecological and reproductive health. Microbiol Mol Biol Rev. 2022;86:e00046–21.

12. Oksanen J, Simpson GL, Blanchet FG, Kindt R, Legendre P, Minchin PR, et al. vegan: Community Ecology Package. R package version 2.6–10. 2025.

13. Fernandes AD, Macklaim JM, Linn TG, Reid G, Gloor GB. ANOVA-like differential expression (ALDEx) analysis for mixed population RNA-seq. PLoS One. 2013;8:e67019.

14. Bastian M, Heymann S, Jacomy M. Gephi: an open source software for exploring and manipulating networks. Proc Int AAAI Conf Web Soc Media. 2009;3:361–362.

15. Parikh HI, Koparde VN, Bradley SP, Buck GA, Sheth NU. MeFiT: merging and filtering tool for Illumina paired-end reads for 16S rRNA amplicon sequencing. BMC Bioinformatics. 2016;17:491.

16. Fettweis JM, Serrano MG, Sheth NU, Mayer CM, Glascock AL, Brooks JP, et al. Species-level classification of the vaginal microbiome. BMC Genomics. 2012;13 Suppl 8:S17.

17. Gravlee CC. Systemic racism, chronic health inequities, and COVID-19: a syndemic in the making? Am J Hum Biol. 2020;32:e23482.

18. Orji AF, Yamashita T. Racial disparities in routine health checkup and adherence to cancer screening guidelines among women in the United States of America. Cancer Causes Control. 2021;32:1247–1256.

