## Supplementary Table 1 for "HPV co-infection burden, vaginal microbiota restructuring, and host context are associated with Cervical precancer risk in a racially restricted observational cohort"

*Participant characteristics in the final Black/African American and White analytic cohort overall and by CIN3 status*

Analytic cohort restricted to participants with non-missing CIN3 status, vaginal microbiome type, and Black/African American/White race classification (N = 1,181).

| **Characteristic** | **Category** | **Overall (N = 1,181)** | **CIN3 (n = 75)** | **No CIN3 (n = 1,106)** | **P value** |
| --- | --- | --- | --- | --- | --- |
| **Race** |  |  |  |  | Pearson χ²(1) = 12.81; P < 0.001 |
|  | White | 359 (30%) | 9 (12%) | 350 (32%) |  |
|  | Black/African American | 822 (70%) | 66 (88%) | 756 (68%) |  |
| **Education** |  |  |  |  | Pearson χ²(4) = 25.15; P < 0.001 |
|  | < High school | 83 (7%) | 6 (8%) | 77 (7%) |  |
|  | High school diploma/GED | 385 (34%) | 42 (58%) | 343 (32%) |  |
|  | Some college | 389 (34%) | 19 (26%) | 370 (35%) |  |
|  | Bachelor's degree | 161 (14%) | 4 (6%) | 157 (15%) |  |
|  | Master's/Doctorate | 119 (10%) | 1 (1%) | 118 (11%) |  |
| **Employment status** |  |  |  |  | Pearson χ²(2) = 14.86; P = 0.001 |
|  | Unemployed/student/homemaker/retired | 490 (45%) | 45 (67%) | 445 (43%) |  |
|  | Part-time | 173 (16%) | 8 (12%) | 165 (16%) |  |
|  | Full-time | 431 (39%) | 14 (21%) | 417 (41%) |  |
| **Annual household income** |  |  |  |  | Pearson χ²(5) = 20.04; P = 0.001 |
|  | < $15,000 | 489 (45%) | 40 (61%) | 449 (44%) |  |
|  | $15,000–$19,999 | 109 (10%) | 12 (18%) | 97 (10%) |  |
|  | $20,000–$39,999 | 194 (18%) | 9 (14%) | 185 (18%) |  |
|  | $40,000–$59,999 | 93 (9%) | 0 (0%) | 93 (9%) |  |
|  | $60,000–$79,999 | 68 (6%) | 3 (5%) | 65 (6%) |  |
|  | ≥ $80,000 | 129 (12%) | 2 (3%) | 127 (12%) |  |
| **Marital status** |  |  |  |  | Pearson χ²(1) = 17.01; P < 0.001 |
|  | Unmarried | 324 (29%) | 5 (7%) | 319 (31%) |  |
|  | Married | 790 (71%) | 64 (93%) | 726 (69%) |  |
| **Smoking status** |  |  |  |  | Pearson χ²(3) = 9.01; P = 0.029 |
|  | Never | 581 (49%) | 29 (39%) | 552 (50%) |  |
|  | Former | 126 (11%) | 14 (19%) | 112 (10%) |  |
|  | Current | 446 (38%) | 32 (43%) | 414 (37%) |  |
|  | Unknown | 28 (2%) | 0 (0%) | 28 (3%) |  |
| **Vaginal microbiome type** |  |  |  |  | Pearson χ²(9) = 15.76; P = 0.072 |
|  | VMB type 1 | 124 (10%) | 13 (17%) | 111 (10%) |  |
|  | VMB type 2 | 243 (21%) | 20 (27%) | 223 (20%) |  |
|  | VMB type 3 | 171 (14%) | 2 (3%) | 169 (15%) |  |
|  | VMB type 4 | 34 (3%) | 3 (4%) | 31 (3%) |  |
|  | VMB type 5 | 342 (29%) | 24 (32%) | 318 (29%) |  |
|  | VMB type 6 | 18 (2%) | 0 (0%) | 18 (2%) |  |
|  | VMB type 7 | 47 (4%) | 2 (3%) | 45 (4%) |  |
|  | VMB type 8 | 52 (4%) | 2 (3%) | 50 (5%) |  |
|  | VMB type 9 | 140 (12%) | 8 (11%) | 132 (12%) |  |
|  | VMB type 10 | 10 (1%) | 1 (1%) | 9 (1%) |  |
| **Any HPV positive (>=25 reads on any type)** |  |  |  |  | Pearson χ²(1) = 34.25; P < 0.001 |
|  | No | 695 (59%) | 20 (27%) | 675 (61%) |  |
|  | Yes | 486 (41%) | 55 (73%) | 431 (39%) |  |
| **Number of HPV types present (>=25 reads)** |  |  |  |  | Pearson χ²(7) = 92.48; P < 0.001 |
|  | 0 | 695 (59%) | 20 (27%) | 675 (61%) |  |
|  | 1 | 275 (23%) | 20 (27%) | 255 (23%) |  |
|  | 2 | 118 (10%) | 13 (17%) | 105 (9%) |  |
|  | 3 | 58 (5%) | 11 (15%) | 47 (4%) |  |
|  | 4 | 21 (2%) | 6 (8%) | 15 (1%) |  |
|  | 5 | 9 (1%) | 3 (4%) | 6 (1%) |  |
|  | 6 | 3 (0%) | 0 (0%) | 3 (0%) |  |
|  | 7 | 2 (0%) | 2 (3%) | 0 (0%) |  |
| **Number of high-risk HPV types present** |  |  |  |  | Pearson χ²(4) = 100.37; P < 0.001 |
|  | 0 | 926 (78%) | 30 (40%) | 896 (81%) |  |
|  | 1 | 205 (17%) | 31 (41%) | 174 (16%) |  |
|  | 2 | 44 (4%) | 10 (13%) | 34 (3%) |  |
|  | 3 | 5 (0%) | 3 (4%) | 2 (0%) |  |
|  | 4 | 1 (0%) | 1 (1%) | 0 (0%) |  |
| **Multiple high-risk HPV infections** |  |  |  |  | Pearson χ²(1) = 41.15; P < 0.001 |
|  | No | 1131 (96%) | 61 (81%) | 1070 (97%) |  |
|  | Yes | 50 (4%) | 14 (19%) | 36 (3%) |  |
| **Metronidazole exposure** |  |  |  |  | Pearson χ²(1) = 0.11; P = 0.743 |
| Metronidazole exposure | No | 851 (84%) | 54 (86%) | 797 (84%) |  |
| Metronidazole exposure | Yes | 159 (16%) | 9 (14%) | 150 (16%) |  |
| **Any antibiotic exposure** |  |  |  |  | Pearson χ²(1) = 2.12; P = 0.146 |
|  | No | 704 (67%) | 49 (75%) | 655 (67%) |  |
|  | Yes | 344 (33%) | 16 (25%) | 328 (33%) |  |
| **Birth control use** |  |  |  |  | Pearson χ²(1) = 3.78; P = 0.052 |
|  | No | 497 (47%) | 38 (58%) | 459 (46%) |  |
|  | Yes | 565 (53%) | 27 (42%) | 538 (54%) |  |
| **Ever pregnant** |  |  |  |  | Pearson χ²(1) = 4.50; P = 0.034 |
|  | No | 958 (85%) | 63 (94%) | 895 (85%) |  |
|  | Yes | 168 (15%) | 4 (6%) | 164 (15%) |  |
| **Prior abnormal Pap smear** |  |  |  |  | Pearson χ²(1) = 1.81; P = 0.178 |
|  | No | 675 (67%) | 35 (59%) | 640 (68%) |  |
|  | Yes | 328 (33%) | 24 (41%) | 304 (32%) |  |
| **Prior colposcopy** |  |  |  |  | Pearson χ²(1) = 23.54; P < 0.001 |
|  | No | 827 (83%) | 38 (60%) | 789 (84%) |  |
|  | Yes | 173 (17%) | 25 (40%) | 148 (16%) |  |
| **Bacterial vaginosis** |  |  |  |  | Pearson χ²(1) = 0.46; P = 0.499 |
|  | No | 672 (67%) | 37 (63%) | 635 (67%) |  |
|  | Yes | 335 (33%) | 22 (37%) | 313 (33%) |  |
| **Yeast infection** |  |  |  |  | Pearson χ²(1) = 1.24; P = 0.266 |
|  | No | 467 (45%) | 33 (52%) | 434 (44%) |  |
|  | Yes | 574 (55%) | 31 (48%) | 543 (56%) |  |
| **Urinary tract infection** |  |  |  |  | Pearson χ²(1) = 1.77; P = 0.184 |
|  | No | 484 (47%) | 34 (55%) | 450 (46%) |  |
|  | Yes | 553 (53%) | 28 (45%) | 525 (54%) |  |
| **Chlamydia** |  |  |  |  | Pearson χ²(1) = 7.46; P = 0.006 |
|  | No | 716 (68%) | 32 (52%) | 684 (69%) |  |
|  | Yes | 333 (32%) | 29 (48%) | 304 (31%) |  |
| **Herpes** |  |  |  |  | Pearson χ²(1) = 0.22; P = 0.637 |
|  | No | 919 (90%) | 53 (88%) | 866 (90%) |  |
|  | Yes | 101 (10%) | 7 (12%) | 94 (10%) |  |
| **Genital warts** |  |  |  |  | Pearson χ²(1) = 10.54; P = 0.001 |
|  | No | 944 (93%) | 48 (83%) | 896 (94%) |  |
|  | Yes | 69 (7%) | 10 (17%) | 59 (6%) |  |
| **Gonorrhea** |  |  |  |  | Pearson χ²(1) = 0.25; P = 0.617 |
|  | No | 855 (81%) | 49 (79%) | 806 (82%) |  |
|  | Yes | 195 (19%) | 13 (21%) | 182 (18%) |  |
| **Syphilis** |  |  |  |  | Pearson χ²(1) = 0.02; P = 0.902 |
|  | No | 975 (98%) | 55 (98%) | 920 (98%) |  |
|  | Yes | 20 (2%) | 1 (2%) | 19 (2%) |  |
| **Trichomonas** |  |  |  |  | Pearson χ²(1) = 1.42; P = 0.233 |
|  | No | 758 (75%) | 40 (69%) | 718 (76%) |  |
|  | Yes | 246 (25%) | 18 (31%) | 228 (24%) |  |
| **Fibroids** |  |  |  |  | Pearson χ²(1) = 0.53; P = 0.466 |
|  | No | 913 (88%) | 57 (90%) | 856 (87%) |  |
|  | Yes | 130 (12%) | 6 (10%) | 124 (13%) |  |
| **Endometriosis** |  |  |  |  | Pearson χ²(1) = 4.66; P = 0.031 |
|  | No | 988 (95%) | 59 (89%) | 929 (95%) |  |
|  | Yes | 52 (5%) | 7 (11%) | 45 (5%) |  |
| **Infertility** |  |  |  |  | Pearson χ²(1) = 2.34; P = 0.126 |
|  | No | 1004 (97%) | 63 (100%) | 941 (96%) |  |
|  | Yes | 35 (3%) | 0 (0%) | 35 (4%) |  |
| **Polycystic ovary syndrome** |  |  |  |  | Pearson χ²(1) = 4.41; P = 0.036 |
|  | No | 839 (86%) | 56 (95%) | 783 (85%) |  |
|  | Yes | 141 (14%) | 3 (5%) | 138 (15%) |  |
| **Pelvic inflammatory disease** |  |  |  |  | Pearson χ²(1) = 0.33; P = 0.567 |
|  | No | 945 (93%) | 57 (95%) | 888 (93%) |  |
|  | Yes | 69 (7%) | 3 (5%) | 66 (7%) |  |
| **Asthma** |  |  |  |  | Pearson χ²(1) = 2.63; P = 0.105 |
|  | No | 829 (78%) | 55 (86%) | 774 (77%) |  |
|  | Yes | 237 (22%) | 9 (14%) | 228 (23%) |  |
| **Diabetes** |  |  |  |  | Pearson χ²(1) = 2.59; P = 0.107 |
|  | No | 981 (94%) | 63 (98%) | 918 (93%) |  |
|  | Yes | 66 (6%) | 1 (2%) | 65 (7%) |  |
| **High blood pressure** |  |  |  |  | Pearson χ²(1) = 1.53; P = 0.216 |
|  | No | 783 (75%) | 44 (69%) | 739 (76%) |  |
|  | Yes | 258 (25%) | 20 (31%) | 238 (24%) |  |
| **High cholesterol** |  |  |  |  | Pearson χ²(1) = 0.51; P = 0.476 |
|  | No | 904 (87%) | 52 (84%) | 852 (87%) |  |
|  | Yes | 137 (13%) | 10 (16%) | 127 (13%) |  |
| **Any cancer history** |  |  |  |  | Pearson χ²(1) = 0.26; P = 0.611 |
|  | No | 1060 (94%) | 67 (93%) | 993 (94%) |  |
|  | Yes | 63 (6%) | 5 (7%) | 58 (6%) |  |

Abbreviations: CIN3, cervical intraepithelial neoplasia grade 3; HPV, human papillomavirus; VMB, vaginal microbiota. Values are presented as n (%). Denominators vary across rows because percentages are based on non-missing responses for each characteristic. P values are from Pearson chi-square tests comparing CIN3-positive and CIN3-negative participants.
