## Supplementary Table 2 for "HPV co-infection burden, vaginal microbiota restructuring, and host context are associated with Cervical precancer risk in a racially restricted observational cohort"

*Variable-level missingness across candidate predictors in the final Black/African American and White analytic cohort*

Analytic cohort restricted to participants with non-missing CIN3 status, vaginal microbiome type, and Black/African American/White race classification (N = 1,181).

| **Domain** | **Characteristic** | **Missing (n)** | **Missing (%)** |
| --- | --- | --- | --- |
| **Gynecologic comorbidity** |  |  |  |
|  | Polycystic ovary syndrome | 201 | 17.0 |
|  | Pelvic inflammatory disease | 167 | 14.1 |
|  | Infertility | 142 | 12.0 |
|  | Endometriosis | 141 | 11.9 |
|  | Fibroids | 138 | 11.7 |
| **Infection history** |  |  |  |
|  | Syphilis | 186 | 15.7 |
|  | Trichomonas | 177 | 15.0 |
|  | Bacterial vaginosis | 174 | 14.7 |
|  | Genital warts | 168 | 14.2 |
|  | Herpes | 161 | 13.6 |
|  | Urinary tract infection | 144 | 12.2 |
|  | Yeast infection | 140 | 11.9 |
|  | Chlamydia | 132 | 11.2 |
|  | Gonorrhea | 131 | 11.1 |
| **Reproductive/gynecologic history** |  |  |  |
|  | Prior colposcopy | 181 | 15.3 |
|  | Prior abnormal Pap smear | 178 | 15.1 |
|  | Birth control use | 119 | 10.1 |
|  | Ever pregnant | 55 | 4.7 |
| **Medication exposure** |  |  |  |
|  | Metronidazole exposure | 171 | 14.5 |
|  | Any antibiotic exposure | 133 | 11.3 |
| **Clinical comorbidity** |  |  |  |
|  | High blood pressure | 140 | 11.9 |
|  | High cholesterol | 140 | 11.9 |
|  | Diabetes | 134 | 11.3 |
|  | Asthma | 115 | 9.7 |
|  | Any cancer history | 58 | 4.9 |
| **Demographic/socioeconomic** |  |  |  |
|  | Annual household income | 99 | 8.4 |
|  | Employment status | 87 | 7.4 |
|  | Marital status | 67 | 5.7 |
|  | Education | 44 | 3.7 |
|  | Smoking status | 0 | 0.0 |
|  | Age | 0 | 0.0 |
|  | Race (Black/African American vs White) | 0 | 0.0 |
| **Microbiome/virologic** |  |  |  |
|  | Vaginal microbiome type | 0 | 0.0 |
|  | Any HPV positive | 0 | 0.0 |
|  | Number of high-risk HPV types | 0 | 0.0 |
|  | Multiple high-risk HPV infections | 0 | 0.0 |
|  | Number of HPV types | 0 | 0.0 |
| **Outcome** |  |  |  |
|  | CIN3 status | 0 | 0.0 |

Missingness percentages are calculated within the final analytic cohort (N = 1,181). Variables included here correspond to candidate predictors considered in descriptive or integrative analyses.
