## Supplementary material for "HPV co-infection burden, vaginal microbiota restructuring, and host context are associated with Cervical precancer risk in a racially restricted observational cohort": STROBE Checklist

*Observational cohort study*

| **Item** | **Section** | **Recommendation** | **Reported on page(s)** |
| --- | --- | --- | --- |
| 1 | Title and abstract | Indicate study design and provide balanced summary | pp. 1-2 |
| 2 | Background/rationale | Scientific background and rationale | p. 3 |
| 3 | Objectives | Specific objectives and hypotheses | p. 3 |
| 4 | Study design | Key elements of study design early in paper | pp. 4-6 |
| 5 | Setting | Setting, locations, dates | p. 4 |
| 6 | Participants | Eligibility and analytic cohort derivation | p. 4 |
| 7 | Variables | Outcomes, exposures, predictors | pp. 4-6 |
| 8 | Data sources/measurement | Measurement and laboratory methods | pp. 4-6 |
| 9 | Bias | Efforts to address bias/limitations | pp. 10-11 |
| 10 | Study size | How study size was arrived at | p. 4 |
| 11 | Quantitative variables | Handling of quantitative variables | pp. 5-6 |
| 12 | Statistical methods | Statistical analyses and modeling | pp. 5-6 |
| 13 | Participants (results) | Numbers at each stage | pp. 6-8 |
| 14 | Descriptive data | Participant characteristics | p. 15; Supplementary Table 1 |
| 15 | Outcome data | Outcome events and summary measures | pp. 6-8 |
| 16 | Main results | Unadjusted and key comparative results | pp. 6-8 |
| 17 | Other analyses | Subgroup, network, and predictive analyses | pp. 7-8 |
| 18 | Key results | Summary of key findings | pp. 9-11 |
| 19 | Limitations | Study limitations | p. 11 |
| 20 | Interpretation | Overall interpretation | pp. 9-11 |
| 21 | Generalisability | External validity/implications | pp. 10-11 |
| 22 | Funding | Funding and role of funders | p. 11 |
